# Programmable Lipid Nanoparticle Targeting via Corona Engineering

**DOI:** 10.64898/2026.02.27.708523

**Authors:** Alisan Kayabolen, Cian Schmitt-Ulms, Alexander Elsener, Francesca Ferraresso, Keira Donnelly, Angela Xinyi Nan, Ian Harris, Samantha Sgrizzi, Aroosa Anwer, Sabrina Pia Nuccio, Patrick T. Paine, Sanah Langer, Christopher Fell, Jonathan S. Gootenberg, Omar O. Abudayyeh

## Abstract

Lipid nanoparticles (LNPs) are a versatile platform for in vivo delivery of biomolecules, yet systemically administered LNPs predominantly accumulate in the liver, limiting extrahepatic applications. This tropism arises from LNP adsorption of serum proteins, particularly apolipoprotein E (ApoE), which binds to LDL receptors (LDLR) on hepatocytes. Here, we overcome this tropism with two compatible strategies. First, we engineer dead ApoE mutants (dApoE) with five receptor-binding domain substitutions that selectively disrupt the ApoE-LDLR interaction but retain lipid binding. In cultured cells, pre-coating with these dApoE markedly inhibited LDLR-mediated uptake. Second, we pretreat cells with hyperactive PCSK9 (haPCSK9) to internalize surface LDLR, similarly reducing the LDLR-mediate uptake of LNPs. In vivo, both strategies substantially reduced liver LNP transduction without inducing redistribution to other major organs. To retarget LNP to new cell types we combined antibody conjugation with dApoE or haPCSK9, effectively engineering tropism to T cells, brain and lung tissues in vivo with substantially reduced hepatic background. In pilot studies, this strategy enabled specific delivery of reporter mRNAs to additional tissues, including megakaryocytes, hematopoietic progenitor cells, and cardiac tissue, and in aged T cells, to deliver miRNA cargos that produced a sustained reduction in DNA damage markers following a single systemic dose. dApoE coated CD5-targeted LNPs generated CAR+ T cells that retained cytotoxicity against CD19+ targets, while simultaneously reducing hepatocyte transduction by 90%. These findings establish a modular framework that integrates dApoE and haPCSK9-mediated detargeting with antibody-based retargeting, allowing for improvements in LNP specificity and broadening the therapeutic scope of LNPs.

## 1. Main

Lipid nanoparticles (LNPs) have emerged as the leading non-viral delivery platform for nucleic acids, enabling the clinical translation of siRNA ^1^ and mRNA therapies ^2^. Their success is rooted in the ability of systemically administered LNPs to accumulate in the liver, where hepatocytes efficiently internalize cargo. However, this tropism represents both a strength and a limitation. For diseases requiring hepatic delivery, ApoE-mediated targeting of LNPs to LDL receptors (LDLR) on hepatocytes is advantageous ^3,4^. Yet, broad application of LNPs to extrahepatic tissues remains restricted by this predominant liver bias.

The mechanism underlying this hepatic tropism is increasingly well understood. After intravenous injection, LNPs rapidly adsorb serum proteins to form a “biomolecular corona” ^5^. Among these proteins, apolipoprotein E (ApoE) plays a dominant role by binding to the LNP surface and facilitating uptake into hepatocytes via LDLR ^6^. Modulating this ApoE-LDLR interaction therefore represents a rational strategy to reduce liver targeting and enable access to other tissues.

Previous approaches to alter LNP tropism include changing ionizable lipid chemistry ^7,8^, or modifying helper or PEG lipids ^9,10^. While effective in part, these methods often lack precision or compatibility with modular targeting approaches. A more selective strategy would be to engineer corona proteins, preserving their lipid-binding capacity while redirecting their receptor-targeting activity.

Beyond detargeting, achieving active retargeting to specific tissues or cell types remains a central challenge. Several groups have explored direct conjugation of ligands, peptides, or antibodies to LNPs, successfully targeting T cells ^11,12^ and hematopoietic stem cells ^13,14^, while others have developed novel lipid formulations (e.g., SORT particles) to passively target the lungs and spleen ^15,16^. While promising, these strategies face limitations in specificity due to hepatic tropism, reproducibility, or in vivo efficiency. An ideal system would combine robust detargeting from the liver with modular retargeting to alternative tissues in a way that is mechanistically clear and broadly applicable.

Here, we report a two-pronged strategy to achieve this goal. First, we blunt liver transduction by engineering both a dead ApoE (dApoE) defective in LDLR binding but competent in lipid binding and a hyperactive PCSK9 with a D377Y mutation (haPCSK9) with increased LDLR surface reduction ^17,18^. We demonstrate that precoating LNPs with dApoE or pretreatment with haPCSK9 effectively reduces hepatic uptake *in vitro* and *in vivo*. Second, we conjugate antibodies to the detargeted LNPs to redirect them toward specific cell or tissue types, including T cells, brain endothelial cells, and lung. Finally, we show the translational potential of this approach by generating functional CAR T cells in vivo using CD5-targeted LNPs combined with liver detargeting strategies. This platform offers a modular framework to expand the therapeutic reach of LNPs beyond the liver while reducing liver transduction.

## 2. Results

### ApoE Mutants Block ApoE-Mediated Uptake and Reduce Liver Accumulation

To directly test whether selective disruption of ApoE-LDLR interactions could prevent liver-specific uptake (**Fig. 1a**), we engineered a series of ApoE mutants carrying substitutions within the receptor-binding domain while preserving the lipid-binding domain. In initial assays, we produced secreted ApoE protein from HEK293FT cells overexpressing either wild-type ApoE (ApoE-WT) or a panel of single, double, triple, and five-residue combinatorial mutants. As expected, pre-coating LNPs with ApoE-WT markedly increased LNP uptake relative to no ApoE control, consistent with its role in binding LNPs to LDLR (**Fig. 1b**). Many single mutants reduced uptake to varying degrees, but combining mutations produced progressively stronger inhibition. The final ApoE construct, combining 5 individual mutations, exhibited the most pronounced blockade of ApoE-mediated enhancement, effectively ablating LNP internalization (**Fig. 1b**). Because this variant showed the strongest and most consistent inhibitory effect, we used it for all subsequent experiments and refer to it hereafter as dead ApoE (dApoE). Quantification of GFP plate-read measurements confirmed the imaging results, showing that while individual mutations partially reduced uptake, the five-mutation ApoE variant (dApoE) produced the strongest and most consistent suppression of ApoE-mediated LNP internalization (**Fig. 1c**).

**Fig. 1.**
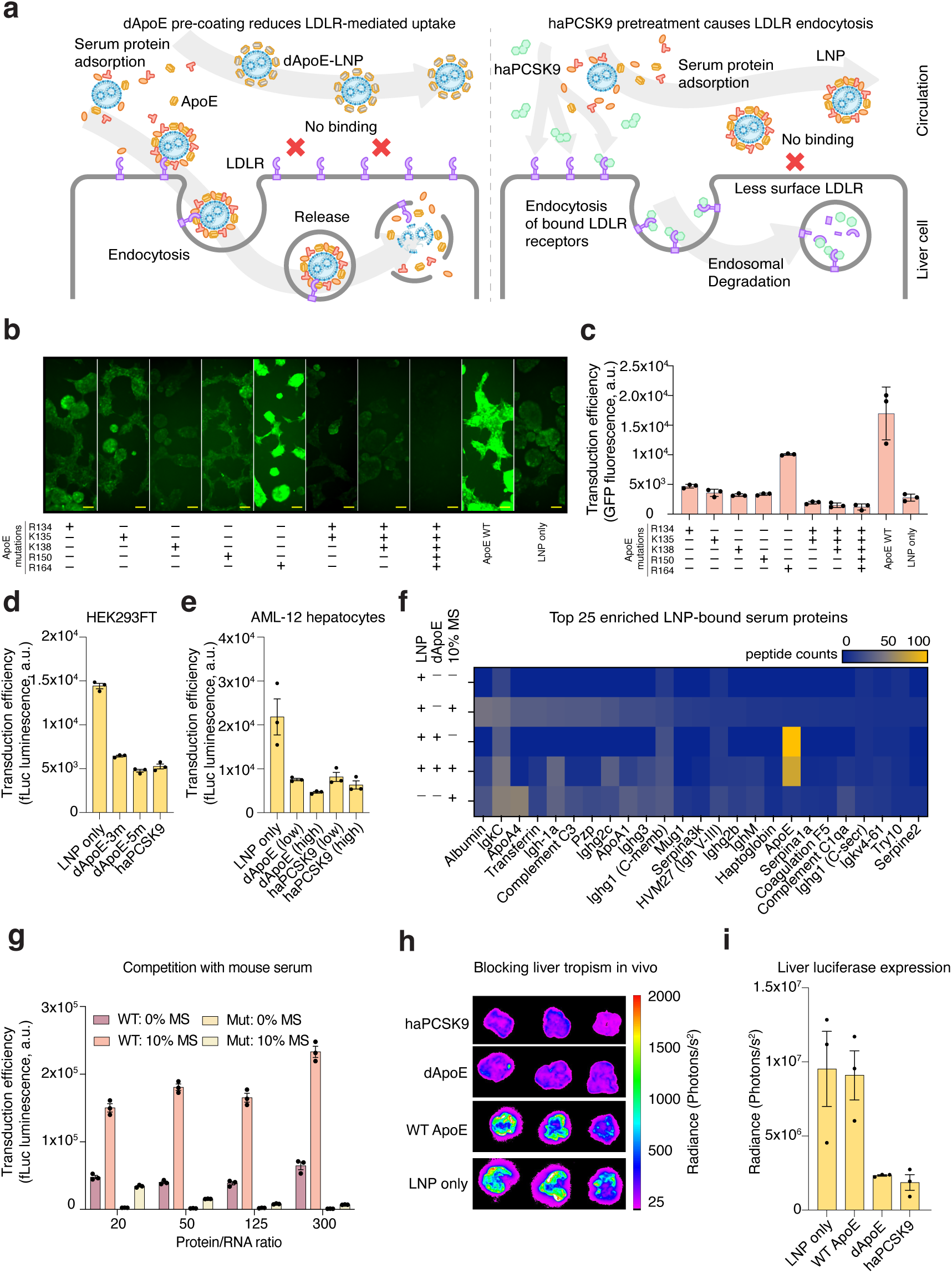
ApoE mutants reduce ApoE-mediated LNP uptake and liver accumulation. (a) Schematic of dApoE precoating and haPCSK9 pretreatment strategies for modulating LNP biodistribution. (b) Representative fluorescence microscopy images showing uptake of GFP-loaded LNPs in HEK293FT cells after incubation with media containing wild-type (WT) ApoE or single, double, triple, and five-residue combinatorial ApoE mutants. Progressive combination of receptor-binding domain mutations resulted in increasingly strong inhibition of ApoE-mediated uptake, with the five-residue mutant exhibiting the most pronounced blockade. Scale bar, 100 µm. (c) Quantification of the LNP uptake by GFP plate-read measurements, confirming that while individual mutations partially reduced uptake, combinatorial mutations produced progressively stronger inhibition, with the five-residue ApoE mutant showing maximal suppression of LNP internalization. (d) Uptake of LNPs in HEK293FT and AML-12 (e) hepatocytes using purified proteins, comparing WT ApoE, three-residue ApoE mutant (dApoE-3m), five-residue ApoE mutant (dApoE-5m), and haPCSK9 pretreatment. Both dApoE and haPCSK9 robustly suppress ApoE-dependent uptake. (f) Proteomic analysis of LNP–protein corona composition showing that dApoE precoating stabilizes the corona and maintains mutant ApoE enrichment even after subsequent serum exposure, whereas LNPs incubated with 10% mouse serum alone recruit a broader range of serum proteins. (g) Serum competition assays demonstrating enhanced uptake with WT ApoE and persistent inhibition by dApoE at higher concentrations, consistent with LC-MS findings on corona stability. (h) In vivo bioluminescence images after 6h of LNP injection (0.3 mg/kg RNA) showing reduced hepatic accumulation with dApoE precoating (25× relative to LNP mRNA) or haPCSK9 pretreatment (40 µg per mouse, administered 15 min before LNP injection). (i) Quantification of bioluminescence across major organs (liver, spleen, kidneys, heart, lungs), showing selective reduction in liver uptake without redistribution to other tissues. All data in this figure are mean ± SEM (n=3).

We next validated the activity of dApoE using purified proteins. In HEK293FT cells, the five-mutation ApoE variant (dApoE) suppressed LNP uptake more effectively than a three-mutation construct, confirming that the inhibitory effect is mutation-dependent and not reliant on other secreted factors present in conditioned media (**Fig. 1d**). Similar inhibition was observed in physiologically relevant AML-12 hepatocytes, demonstrating that the dApoE effect was not cell line specific (**Fig. 1e**). As an orthogonal detargeting strategy, we applied haPCSK9 pretreatment, which induces LDLR degradation. haPCSK9 phenocopied the effect of dApoE across both cell types, validating LDLR as the primary mediator and offering an alternative, therapeutically relevant mechanism for modulating hepatic uptake (**Fig. 1d-e**).

To characterize how ApoE mutations alter corona composition and simulate an *in vivo* environment, we performed LC-MS proteomic profiling of LNPs after incubation with either dApoE, 10% mouse serum (MS10), or dApoE followed by MS10 exposure. dApoE precoating stabilized the corona and maintained mutant ApoE enrichment even after subsequent serum exposure, whereas serum-only samples recruited a broader range of proteins (**Fig. 1f**). These data indicate that dApoE not only blocks LDLR-mediated uptake but also reshapes the corona to favor its own retention over exchange with serum components.

Competition assays performed in the presence of serum further confirmed specificity: ApoE-WT consistently enhanced LNP uptake, while dApoE retained inhibitory effects even at low concentrations (**Fig. 1g**). Notably, these functional results were consistent with the LC-MS findings (**Fig. 1f**), which showed that dApoE precoating stabilizes its association with LNPs and limits exchange with serum proteins.

We next evaluated these strategies *in vivo*. Both dApoE precoating and haPCSK9 pretreatment markedly reduced hepatic accumulation following systemic administration (**Fig. 1h-i**). Importantly, analysis of all major organs showed that detargeting did not lead to redistribution into unintended tissues (**Extended Data Fig. 1a**). Signals in spleen, kidneys, heart, and lungs were comparable across groups, indicating that reducing liver uptake does not simply shift accumulation to other sites. Rather, detargeting selectively suppresses hepatic uptake without causing nonspecific redistribution. Consistent with reduced hepatic sequestration, detargeting with dApoE or PCSK9 increased circulating fLuc mRNA levels in blood, indicating prolonged LNP persistence in the systemic compartment as a result of reduced liver uptake (**Extended Data Fig. 1b**). A pilot experiment showed that dApoE mediated liver detargeting is dose-dependent (**Extended Data Fig. 1c-d**), and additional lipid-binding domain mutations have the potential to enhance LNP binding and in vivo liver detargeting (**Extended Data Fig. 1e-f**). Collectively, these data establish that dApoE and haPCSK9 act through a shared LDLR-dependent mechanism, and that detargeting can be achieved without compromising extrahepatic delivery potential.

### Modular Retargeting via Fusion Proteins and Antibody Conjugation

Having established a robust strategy for detargeting the liver, we next explored whether detargeted LNPs could be actively redirected to alternative cell types (**Fig. 2a**). As a first step, we tested modular fusions in which synthetic binders (**Extended Data Fig. 2a-b**) or single-chain variable fragments (scFvs) (**Extended Data Fig. 2c-d**) were genetically linked to dApoE. These constructs enabled receptor-specific uptake, but only in cells engineered to overexpress the corresponding receptors, and the magnitude of uptake enhancement was relatively modest. While this confirmed the principle of modular retargeting, the efficiency achieved was insufficient for translational applications.

**Fig. 2.**
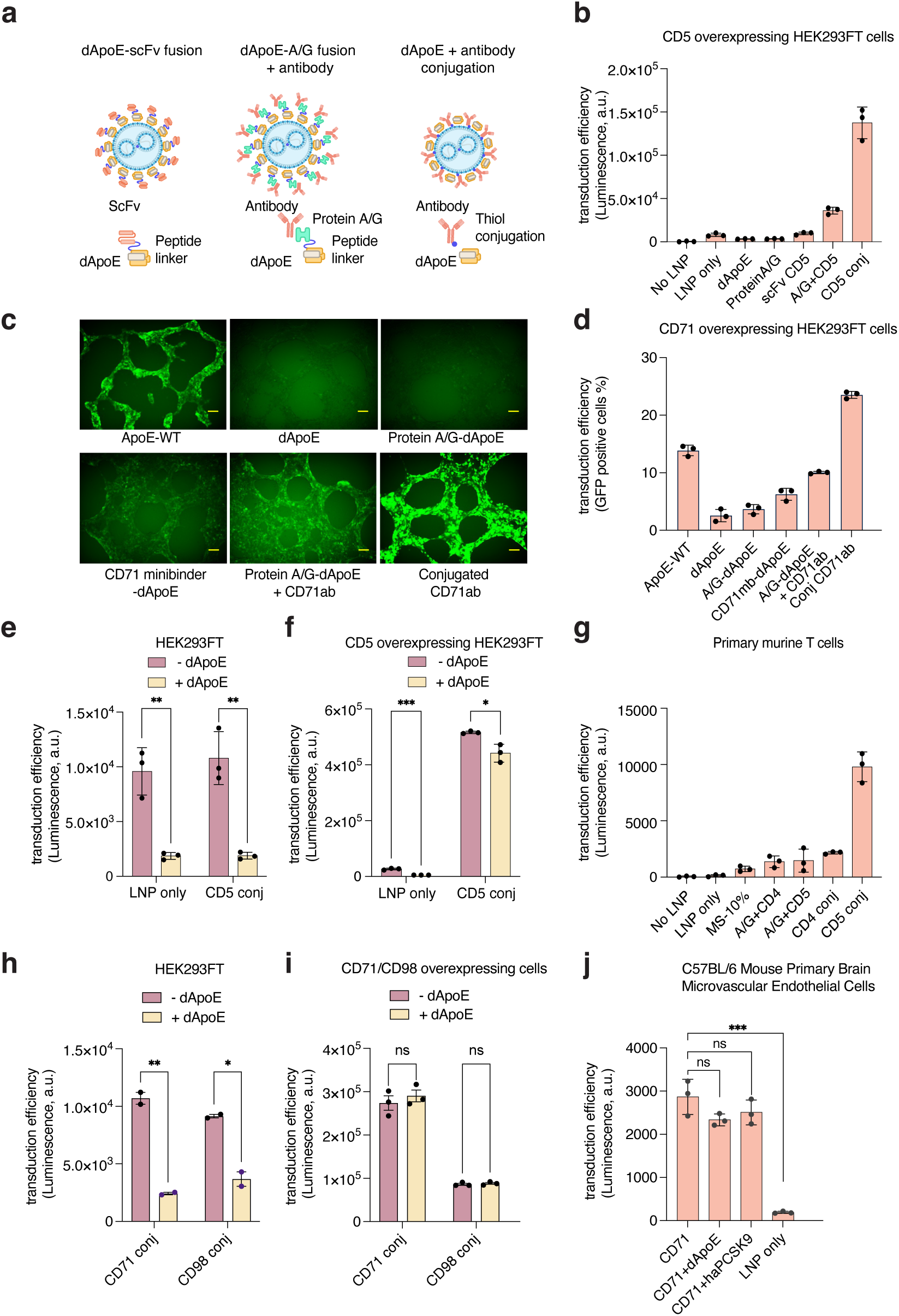
Retargeting LNPs via fusion proteins and antibody conjugation. (A) Schematic of retargeting strategies: dApoE-ligand fusion constructs, dApoE-Protein A/G fusion followed by antibody incubation, and direct antibody conjugation. (B) Comparison of 3 different retargeting strategies in CD5-overexpressing HEK293FT cells. Precoating fLuc-LNPs with secreted dApoE or dApoE-Protein A/G fusion inhibits cellular uptake, while fusion with either CD5-targeting scFv or Protein A/G followed by CD5 antibody addition improve uptake efficiency. However, the strongest signal is obtained by direct conjugation of CD5 antibodies on LNPs. (C) Fluorescence imaging of CD71-targeting GFP-LNPs in transferrin receptor (CD71)-overexpressing HEK293FT cells at 24h. dApoE or dApoE-Protein A/G precoating inhibits cellular uptake of LNPs. CD71 minibinder-dApoE fusions and Protein A/G-dApoE plus CD71 antibody produce modest uptake, whereas direct CD71 antibody conjugation yields the strongest and most uniform signal. Scale bar, 100 µm. (D) Flow cytometry analysis of GFP uptake at 72h in HEK293FT cells for CD71 targeting, with antibody conjugation compared to minibinder or Protein A/G fusion strategies. (E) Luminescence-based quantification of CD5-targeted LNP uptake in HEK293FT cells; effect of dApoE precoating on background signal. (F) Transduction efficiency of CD5 antibody-conjugated and unconjugated LNPs in CD5-overexpressing HEK293FT cells, showing sustained targeting and reduced off-target uptake with dApoE. (G) Transduction in primary murine T cells using Protein A/G-dApoE plus antibody, or directly conjugated CD4/CD5 antibodies; CD5 conjugation yields the strongest signal. (H) Luminescence-based quantification of CD71- or CD98-targeted LNP uptake in HEK293FT cells; effect of dApoE precoating on background signal. (I) Transduction efficiency of CD71 or CD98 antibody-conjugated LNPs in CD71- or CD98-overexpressing HEK293FT cells, showing sustained targeting with dApoE. (J) Enhanced delivery to primary brain endothelial cells using CD71 antibody-conjugated LNPs; unaffected by dApoE or haPCSK9 detargeting. All data in this figure are mean ± SEM (n=3).

We then compared three retargeting strategies directly: 1) scFv, binder or ligand fusions to dApoE, 2) dApoE fused to Protein A/G followed by antibody addition, and 3) direct chemical antibody conjugation. Using CD5-overexpressing HEK293FT cells and fLuc-LNPs, we found that antibody-conjugated LNPs produced the most robust receptor-specific uptake, whereas fusion-based approaches resulted in substantially lower enhancement (**Fig. 2b**). To determine whether this hierarchy generalizes across receptor systems, we performed parallel experiments targeting the transferrin receptor (CD71). Here, we compared GFP-loaded LNPs precoated either with dApoE fused with CD71 minibinder, or Protein A/G-dApoE fusion plus CD71 antibody, to CD71 antibody-conjugated LNPs. Fluorescence imaging at 24h revealed detectable enhancement in the uptake with both fusion-based strategies, but again the brightest and most uniform delivery was achieved with antibody-conjugated LNPs (**Fig. 2c**). Quantitative flow cytometry at 72 h confirmed these results, showing that conjugated CD71-LNPs outperformed all fusion formats (**Fig. 2d**). These results are consistent with the CD5 dataset and indicate that direct antibody conjugation is the most efficient of the tested retargeting modalities in combination with dApoE coronas. In subsequent experiments, we tested to combine antibody-mediated targeting strategy with dApoE-mediated detargeting. First, we compared CD5-targeted transduction in plain and CD5-overexpressing HEK293FT cells in the presence or absence of purified dApoE. In receptor-negative HEK293FT cells (**Fig. 2e**), both unconjugated and CD5-conjugated LNPs yielded similar off-target luminescence signals, but pre-coating with dApoE efficiently blocked nearly 80–85% of this background uptake. In CD5-overexpressing cells (**Fig. 2f**), CD5 antibody-conjugated LNPs demonstrated roughly 200-fold higher transduction relative to unconjugated LNPs, and importantly, dApoE pre-treatment suppressed only the off-target uptake without diminishing the specific CD5-mediated delivery.

Next, we evaluated whether these findings generalize to primary immune cells by comparing targeting strategies in primary murine T cells (**Fig. 2g**). Both Protein A/G-dApoE plus antibody incubation and direct antibody conjugation were tested using CD4 and CD5 antibodies. While CD5 targeting outperformed CD4 in both formats, direct antibody conjugation yielded substantially higher transduction than the fusion-based approach for both antibodies. The strongest delivery was observed with CD5 antibody-conjugated LNPs, confirming robust receptor-specific targeting in primary T cells.

To assess whether these findings generalize across receptor systems, we evaluated CD71 antibody-conjugated LNPs in both CD71-overexpressing and parental HEK293FT cells, with or without dApoE pre-coating (**Fig. 2h**). CD71 conjugation enhanced transduction efficiency by 250- to 1000-fold in receptor-positive cells, and importantly, this enhancement was preserved in the presence of dApoE (**Fig. 2i**. In contrast, dApoE pre-coating consistently suppressed background uptake in receptor-negative cells, confirming that detargeting selectively reduces non-specific internalization without compromising receptor-mediated delivery.

A similar pattern was seen for CD98 antibody-conjugated LNPs (**Fig. 2h-i**). In CD98-overexpressing HEK293FT cells, transduction was increased 100-250 fold by direct antibody conjugation and remained high with dApoE treatment, whereas background in receptor-negative cells was efficiently blocked.

Finally, CD71 antibody-conjugated LNPs were tested on C57BL/6 Mouse Primary Brain Microvascular Endothelial Cells (**Fig. 2j**). Here, direct conjugation facilitated a ∼15-fold increase in transduction compared to unconjugated LNPs, and both dApoE pre-coating and haPCSK9 pretreatment maintained the targeted signal without introducing off-target uptake.

### *In Vivo* Targeting of T Cells, Brain, and Lung

Having established that antibody conjugation provides robust and modular retargeting in vitro in combination with dApoE coronas, we next evaluated our strategy *in vivo* across multiple extrahepatic tissues. For these studies, we used firefly luciferase (fLuc) mRNA as a reporter payload to enable quantitative imaging of tissue-specific expression (**Fig. 3a**).

**Fig. 3.**
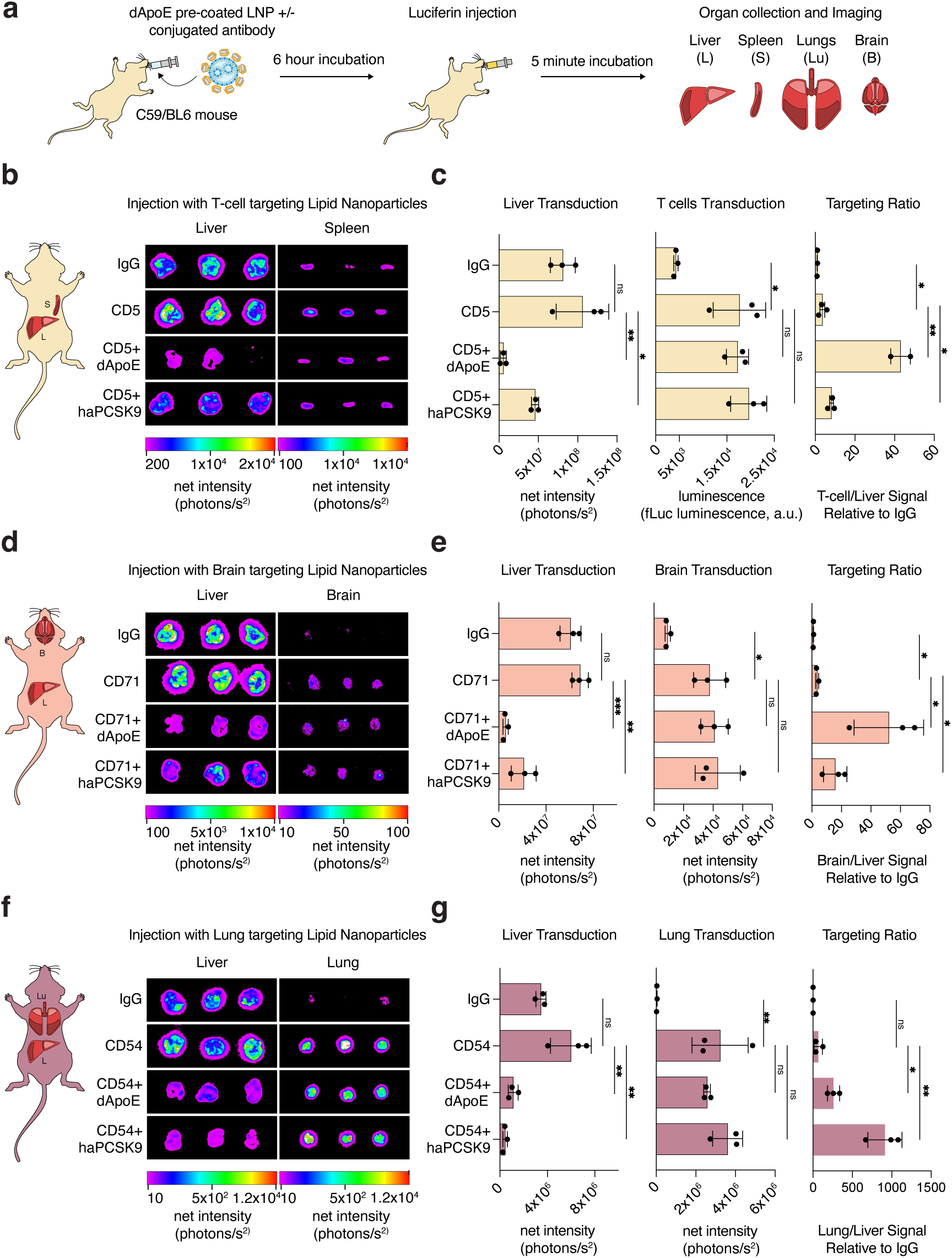
In vivo targeting of T cells, brain, and lung using antibody-conjugated LNPs. (A) Schematic of in vivo targeting experiments. (B) Bioluminescence images showing T cell targeting with CD5 or IgG antibody-conjugated LNPs (0.5 mg/kg RNA) compared to controls, with or without dApoE precoating (25× relative to LNP mRNA) or haPCSK9 pretreatment (40 µg/mouse, 15 min before LNP injection). Quantification of T cell and liver signals, and T cell-to-liver ratios, demonstrating that antibody conjugation increases T-cell delivery while detargeting strategies reduce hepatic background without impairing targeting. (D) Bioluminescence images showing brain-associated uptake of CD71 or IgG antibody-conjugated LNPs (0.5 mg/kg RNA) under control or detargeting conditions. (E) Quantification of brain and liver signals, and brain-to-liver ratios, highlighting improved targeting specificity when combined with dApoE or haPCSK9. (F) Bioluminescence images of liver and lung following systemic administration of CD54 or IgG antibody-conjugated LNPs (0.5 mg/kg RNA). CD54 conjugation results in robust lung delivery, and combining CD54 targeting with dApoE precoating (25× relative to LNP mRNA) or haPCSK9 pretreatment (40 µg per mouse, administered 15 min before LNP injection) further suppresses hepatic accumulation while preserving lung uptake. (G) Quantification of lung and liver signals, and lung-to-liver ratios, demonstrating that CD54 conjugation enables selective lung targeting when combined with detargeting strategies. All data in this figure are mean ± SEM (n=3).

We began by assessing T cell delivery using CD5 antibody-conjugated LNPs, as CD5 is a pan-T cell surface marker broadly expressed across naive and activated T cell subsets and has been previously leveraged for selective targeting and manipulation of T cells in vivo ^12,19,20^. CD5 antibody-conjugated LNPs markedly enhanced uptake into T cells compared to IgG-conjugated controls (**Fig. 3b-c**). Importantly, liver accumulation was substantially reduced by either dApoE precoating or haPCSK9 pretreatment, yet T cell delivery remained unaffected. As a result, the T cell-to-liver signal ratio was significantly increased in detargeted groups, highlighting the ability of this combined strategy to improve both selectivity and efficiency.

We next assessed brain targeting using CD71 antibody-conjugated LNPs, as CD71 (transferrin receptor) is highly expressed on brain endothelial cells and has been extensively exploited to enable receptor-mediated transport across the blood-brain barrier for biologics and nanoparticle-based delivery systems ^21–23^. Similarly, conjugation of CD71 antibodies enhanced uptake into the brain (**Fig. 3d-e**). While IgG controls yielded low background signals, CD71 conjugates produced robust brain expression, and liver detargeting with dApoE or haPCSK9 further increased the brain-to-liver signal ratio. These findings indicate that antibody-mediated targeting enhances uptake in brain-associated cells, likely including endothelial populations, and that detargeting strategies further improve brain-to-liver signal ratios by minimizing hepatic sequestration.

To expand the platform beyond immune and brain-associated compartments, we next assessed lung delivery using CD54 (ICAM-1) antibody-conjugated LNPs, as ICAM-1 is broadly expressed across multiple lung cell populations; including endothelial, epithelial, and immune cells, as shown by single-cell atlases, and has been widely targeted for lung-directed delivery in models of pulmonary inflammation, infection, and cancer ^24–29^. CD54 conjugation resulted in robust lung uptake, far exceeding background observed with IgG controls (**Fig. 3f**). When combined with dApoE precoating or haPCSK9 pretreatment, lung delivery was maintained while liver signal was markedly reduced, yielding substantial increases in the lung-to-liver ratio (**Fig. 3g**). These results establish CD54 conjugation as a strong lung-targeting modality and demonstrate that ApoE-based detargeting synergizes with antibody-mediated delivery to further enhance specificity.

Pilot experiments exploring optimized antibody-to-LNP ratios and dApoE or haPCSK9 dosing revealed opportunities for further enhancement. CD5-conjugated LNPs showed improved T cell targeting efficiency when conjugation and detargeting conditions were adjusted (**Extended Data Fig. 3a-d**), while CD71-conjugated LNPs exhibited higher brain-to-liver signal ratios with optimized dosing regimens (**Extended Data Fig. 3e-h**). These findings suggest that fine-tuning conjugation chemistry, detargeting conditions and dosing parameters can further enhance the efficacy of the combined strategy.

We also extended this approach to additional therapeutically relevant targets. CD98 antibody conjugation increased uptake in brain-associated cells, with haPCSK9 pretreatment further improving brain-to-liver ratios (**Extended Data Fig. 4a**). CD41 conjugation enabled delivery to megakaryocytes, an effect that was maintained or enhanced when combined with dApoE precoating **(Extended Data Fig. 4b**). CD117 antibody conjugation targeted hematopoietic progenitor cells (HPCs), and haPCSK9 pretreatment reduced liver uptake without compromising, and in some cases improving, HPC delivery **(Extended Data Fig. 4c)**. Finally, Nav1.5 (SCN5A) antibody conjugation enhanced cardiac delivery, which was further improved with haPCSK9 pretreatment (**Extended Data Fig. 4d)**. These results mirror the trends observed for T cell, brain, and lung targeting: antibody conjugation provides efficient delivery to target cells, while detargeting strategies consistently lower hepatic background and can even augment on-target accumulation.

### *In Vivo* CAR T Cell Generation

To evaluate the translational potential of our platform, and to assess whether our detargeting-retargeting framework could be applied to generate functional CAR T cells *in vivo*, we formulated LNPs carrying CD19 CAR mRNA and administered them systemically (**Fig. 4a**). After six hours, spleen- and liver-derived cells were isolated for imaging, flow cytometry, and functional assays.

**Fig. 4.**
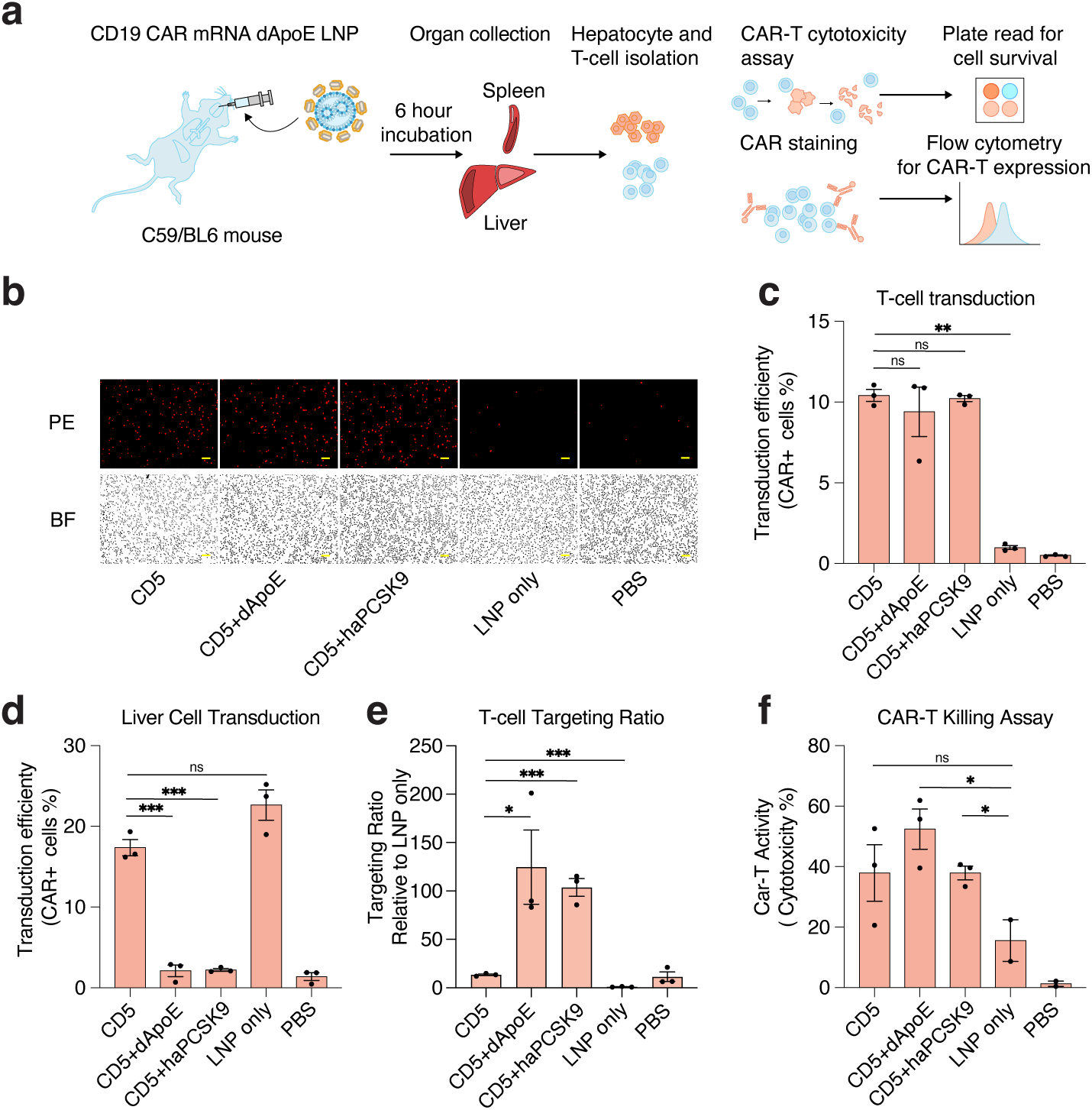
In vivo generation of functional CAR T cells using detargeted and retargeted LNPs. (A) Schematic of CAR T cell generation strategy. (B) Representative microscope images showing CAR expression in T cells isolated from mice treated with CD5-conjugated or unconjugated LNPs (0.5 mg/kg RNA) with or without dApoE precoating (25× relative to LNP mRNA) or haPCSK9 pretreatment (40 µg per mouse, administered 15 min before LNP injection). Scale bar, 50 µm. (C) Flow cytometry quantification of CAR+ T cells following treatment with CD5-conjugated or unconjugated LNPs. (D) CAR expression in hepatocytes from treated animals, showing reduced liver CAR+ cells with detargeting strategies. (E) T-cell targeting ratio showing the ratio of T-cell transduction over liver transduction, normalized to LNP only (F) Functional killing assay showing cytotoxicity of engineered CAR T cells against CD19- and fLuc-overexpressing HEK293FT cells at 24h of co-culture. Cytotoxicity is calculated as the percent decrease in the luminescence signal compared to control. All data in this figure are mean ± SEM (n=3).

Direct fluorescence microscopy revealed robust CAR expression in splenic T cells treated with CD5 antibody-conjugated LNPs, whereas minimal signal was observed with unconjugated LNPs or PBS controls (**Fig. 4b**). Precoating LNPs with dApoE or pretreating animals with haPCSK9 did not diminish CAR expression in T cells, indicating that detargeting does not interfere with on-target transduction. Flow cytometry confirmed these observations quantitatively (**Fig. 4c**). CD5-conjugated LNPs generated approximately 8-12% CAR+ T cells across experiments, compared to ∼1% with unconjugated LNPs and negligible levels with PBS. dApoE or haPCSK9 treatment preserved this efficiency, demonstrating compatibility between the detargeting mechanism and CD5-mediated targeting. In contrast, liver transduction was strongly reduced by detargeting strategies (**Fig. 4d**). In the absence of detargeting, ∼20% of hepatocytes were CAR+, whereas precoating with dApoE or pretreatment with haPCSK9 reduced this to ∼2-3%. Thus, ApoE-LDLR disruption sharply suppresses hepatic off-target editing while maintaining robust T cell delivery.

To evaluate the functional competence of in vivo-generated CAR T cells, we performed a short-term cytotoxicity assay using CD19+ target cells (**Fig. 4e**). CAR T cells derived from mice treated with CD5-conjugated LNPs, with or without dApoE or haPCSK9, exhibited substantial killing activity (40–60% target cell death), whereas control groups showed minimal cytotoxicity. These findings demonstrate that CAR expression induced in vivo is not only efficient but also functionally productive, and that combining targeted delivery with dApoE or haPCSK9-mediated detargeting does not impair T cell effector function.

Together, these data demonstrate that combining antibody-mediated targeting with dApoE or haPCSK9-mediated detargeting enables selective and functional in vivo generation of CAR T cells. Notably, pilot optimization studies exploring higher LNP doses (1 mg/kg RNA) and increased haPCSK9 pretreatment (80 µg per mouse) achieved CAR+ T cell frequencies of ∼35% **(Extended Data Fig. 5a-c)**, suggesting that refined formulation and dosing parameters can further enhance the efficiency of this approach.

### *In vivo* partial reprogramming of aged T cells

To explore whether our targeted-detargeted LNP delivery could support functional rejuvenation of aged immune cells, we performed a pilot experiment that, to our knowledge, represents the first demonstration of T cell-targeted in vivo partial reprogramming in aged mice using miRNA mimics from the previously described miR-302/367 cluster.^30,31^. (**Extended Data Fig. 5d**). Following *in vivo* delivery of CD5-conjugated, haPCSK9-detargeted LNPs carrying partial reprogramming miRNA mimics, γH2AX levels in aged T cells were transiently increased at day 1 relative to age-matched untreated controls. This result is consistent with transcriptional stress-associated DNA damage during reprogramming initiation ^32,33^. By day 5, γH2AX signal was significantly reduced in treated cells compared to controls, indicating resolution of DNA damage over time. This reduction persisted through day 12, where γH2AX levels remained significantly lower in the treated group, although the magnitude of the difference was partially attenuated relative to day 5. These data demonstrate that a single, targeted LNP dose of 302/367 miR mimics can induce a significant reduction in DNA damage markers in aged T cells.

## 3. Discussion

Our study establishes a modular framework that integrates mutant ApoE- or haPCSK9-mediated liver detargeting with antibody-based retargeting to enable precise and programmable LNP delivery beyond the liver. By engineering ApoE mutants with impaired LDLR binding but preserved lipid association, and by leveraging haPCSK9-mediated transient LDLR downregulation, we demonstrate that the ApoE-LDLR axis can be selectively disrupted without compromising the fundamental physicochemical properties that govern LNP performance. Both approaches robustly reduced liver uptake across multiple *in vitro* and *in vivo* settings, underscoring the central role of ApoE-driven hepatocyte internalization in dictating systemic LNP biodistribution.

Importantly, ApoE detargeting did not lead to significant redistribution into unintended tissues. Instead, both dApoE and haPCSK9 selectively suppressed hepatic uptake while maintaining low background in other organs. This property is particularly advantageous compared to approaches relying on lipid chemical modifications, which may unpredictably alter circulation time and biodistribution. The clean detargeting profile observed here suggests that ApoE disruption can serve as a generalizable “background suppression” strategy that improves the specificity and safety margins of a wide range of LNP-based therapeutics.

We further show that liver detargeting preserves, but does not intrinsically increase, antibody-mediated retargeting. Across CD5, CD71, and CD54 conjugates, on-target uptake remained similar with or without detargeting, whereas hepatic background was consistently reduced by dApoE precoating or haPCSK9 pretreatment. As a result, apparent targeting specificity improved substantially, driven primarily by decreases in off-target liver uptake rather than increases in delivery to the intended tissue. Although our brain and lung studies do not distinguish endothelial from parenchymal uptake, they clearly demonstrate that detargeting sharpened the contrast between target and non-target tissues. Thus, ApoE-pathway disruption acts as a generalizable background-suppression strategy that enhances the specificity, rather than the absolute magnitude, of antibody-guided targeting.

Several factors may contribute to why on-target signals remained similar and did not increase with our detargeting strategies. First, even when hepatocyte uptake is inhibited, intravenously dosed LNPs may still physiologically pool in high-capacity reticuloendothelial system organs (liver, spleen); particles may accumulate or be sequestered extracellularly/sinusoidally, limiting the fraction reaching target beds ^34–36^. Second, target-cell receptor availability likely saturates at the tested antibody densities/doses; once occupancy is near maximal in control conditions, further detargeting cannot raise on-target binding. Third, hemodynamic and barrier constraints (e.g., endothelial filtration, interstitial transport) and rapid mononuclear phagocyte system clearance can cap delivery to extrahepatic tissues. Fourth, our readouts quantify protein expression (luminescence/flow), not RNA exposure; reductions in off-target translation do not necessarily imply more RNA translation in on-target cells, and endosomal escape and translational efficiency may remain rate-limiting.

Besides antibody conjugation, ligand- and scFv-fusion approaches also produced receptor-dependent uptake but with lower overall efficiency. A likely contributing factor is that these fusions interact with LNPs through a surface corona rather than covalent attachment. Even though dApoE-mediated corona formation is relatively stable, it remains dynamic and may offer less controlled ligand orientation, valency, and spatial density compared to chemical conjugation. These biophysical constraints could limit avidity for cell-surface receptors, explaining why fusion-based constructs achieve specificity but not the robust uptake observed with covalent antibody conjugation. In contrast, direct antibody-LNP conjugation ensures stable ligand presentation and high functional affinity, consistent with the stronger and more reproducible retargeting effects seen across all evaluated target receptors.

The ability to combine detargeting with antibody-based retargeting creates opportunities for new therapeutic applications. As a translational demonstration, we applied CD5-conjugated, detargeted LNPs to generate CAR T cells in vivo. This approach yielded ∼10-35% CAR+ T cells, an order-of-magnitude improvement over unconjugated LNPs. Detargeting reduced hepatic CAR expression by ∼10-fold, a meaningful safety benefit for systemic CAR delivery. The CAR T cells produced in vivo retained potent cytotoxicity against CD19+ targets, confirming that mRNA delivery and subsequent receptor expression were functional. These results establish a proof-of-concept for in vivo CAR T cell generation using detargeted, antibody-conjugated LNPs, with further optimization of formulation and dosing likely to enhance efficiency toward clinically relevant levels.Beyond T cells, our pilot studies demonstrate that this framework can be extended to additional therapeutically relevant targets. CD71 and CD98 antibody conjugation increased delivery to brain-associated cells; CD41 antibody conjugation enabled megakaryocyte targeting; CD117 antibody conjugation targeted hematopoietic progenitor/stem cells; and Nav1.5 antibody conjugation improved cardiac delivery. In each case, detargeting strategies maintained or further improved specificity. These findings collectively support the generality of combining ApoE detargeting with antibody-mediated retargeting as a modular system for diverse cell types, with further optimization of dosing and conjugation parameters likely to enhance performance across these applications.

Finally, we tested whether T cell-targeted and liver-detargeted LNPs could be leveraged to deliver miRNA mimics and drive tissue specific partial reprogramming *in vivo*. For this proof-of-concept, we selected mimics of the miRNA 302/367 cluster previously shown to induce cellular reprogramming and ameliorate aging hallmarks^30,31^. Using CD5 antibody-conjugated LNPs combined with haPCSK9-mediated liver detargeting, a single systemic dose of miR-302/367 mimics, produced an initial transient increase in DNA damage markers followed by a sustained reduction at later time points. This biphasic response is consistent with early reprogramming-associated stress and subsequent amelioration mediated by upregulation of DNA damage repair pathways^32,33,37^. Although limited in scale, to our knowledge, this is the first demonstration of targeted in vivo, tissue specific rejuvenation via partial reprogramming. Critically, the combination of inherently transient and short-lived miRNA species in a hepatic exempt, tissue specific LNP addresses all key translational hurdles that currently plague the reprogramming field, notably liver toxicity, stochastic off-target reprogramming, and over reprogramming with associated loss of function or even teratoma formation^38,39^. Ultimately, further in vivo studies incorporating other tissue/cell types, expanded cohorts, and functional readouts with optimized dosing are required to define the translational potential of this strategy for the clinical treatment of age associated diseases.

Our study has several limitations. First, the structural basis for the altered LDLR binding of dApoE variants remains to be fully defined, and future biophysical studies will help refine the design principles for next-generation ApoE mutants. Second, while antibody conjugation is effective, large-scale manufacturing and regulatory considerations will require standardized and scalable chemistries. Third, although our pilot experiments demonstrate feasibility across multiple targets, parenchymal penetration, particularly in solid tissues such as brain, lung, and heart, may require additional advances in barrier modulation or ligand discovery. Addressing these challenges will be essential for realizing the full therapeutic potential of the platform.

Our results establish a unified strategy in which dApoE or haPCSK9-mediated detargeting minimizes hepatic sequestration while antibody conjugation drives tissue-specific uptake. Together, these approaches enable flexible, high-specificity LNP delivery with broad potential applications in immunotherapy, neurology, regenerative medicine, and beyond.

In summary, we present a modular platform that combines dApoE or haPCSK9-mediated detargeting with antibody-mediated retargeting to enable selective and efficient LNP delivery beyond the liver. Our results demonstrate the feasibility of this strategy for in vivo CAR T cell generation and suggest broad applicability to brain, lung, cardiac, and hematopoietic targets, among others. By reducing hepatic off-target uptake while preserving on-target delivery, this approach has the potential to significantly expand the therapeutic reach of LNP technologies and improve their safety profile for systemic applications.

## Methods

### LNP Formulation and Characterization

LNPs were formulated by combining ALC-0315 (MedChemExpress) as ionizable cationic lipid, DSPC (Avanti Research) as helper lipid, cholesterol (Avanti Research), and DMG-PEG 2000 (Avanti Research) at a molar ratio of 50:10:38.5:1.5. Lipids were mixed with RNA solution at a 1:3 volume ratio, and at an N:P ratio of 6. For initial in vitro optimization experiments, LNPs were produced by vortex method ^40^. For large scale in vivo experiments, a T-tube setup with dual Harvard Apparatus syringe pump was used ^41^.

Firefly Luciferase (fLuc) or GFP mRNAs were purchased from Terrain Bio or TriLink Biotechnologies, respectively, and used as reporter mRNAs. CD19 CAR mRNA was purchased from MedChemExpress and used to produce CAR-T cells generating LNPs. Average size, size distribution, and polydispersity index (PDI) of the LNPs were measured using Zetasizer Dynamic Light Scattering (DLS) instruments. RNA encapsulation efficiency was measured using the Thermo Fisher RiboGreen assay to quantify RNA. Briefly, diluted LNP samples were treated either with Tris-EDTA (TE) only or Tris-EDTA + detergent, 1% Triton-X, (TEx) and free RNAs were detected using an RNA-binding fluorescent Ribogreen dye. Encapsulated RNA was calculated by subtracting unencapsulated RNA from total RNA, by measuring fluorescence of TE or TEx-treated samples, respectively, in a plate reader and quantified by generating a standard curve using RNA standards.

### Antibody Conjugation on LNPs

To prepare antibody-conjugated LNPs, we utilized SATA-maleimide conjugation chemistry ^42^. LNPs were prepared by incorporating DSPE-PEG-maleimide during the initial lipid formulation process. The original lipid ratio was modified as 50:10:38:1.5:0.5, for ionizable lipid, DSPC, cholesterol, DMG-PEG, and DSPE-PEG-maleimide, respectively.

First, targeting antibodies and control isotype-matched IgG were functionalized with N-succinimidyl S-acetylthioacetate (SATA). SATA was added to the antibodies in a 10-fold molar excess and incubated for 30 minutes at room temperature, introducing sulfhydryl groups. The SATA-modified antibodies were then deprotected using 0.5 M hydroxylamine for 2 hours at room temperature. Unreacted components were removed using Zeba Spin Desalting Columns (7K MWCO, Thermo Scientific) after both steps. The reactive sulfhydryl groups on the antibodies were then conjugated to the maleimide moieties on the LNPs using thioether conjugation chemistry, performed simply by mixing functionalized antibodies with LNPs at different ratios, and incubating at room temperature for 1h to ensure efficient binding.

Following conjugation, the antibody-LNP complexes were purified by qEV size exclusion chromatography columns (70 nm, Izon Science) to remove unbound antibodies and other reaction by-products. LNP-bound proteins were quantified by CBQCA assay (Invitrogen) using the manufacturer’s protocol. Final mRNA content within the LNPs were quantified using Quant-iT RiboGreen RNA assay (Invitrogen). After conjugation, all targeted and non-targeted LNP preparations were stored at 4°C and used within three days of preparation.

### Cell Culture

HEK293FT and AML-12 cells were purchased from ATCC. HEK293FT cells were cultured in DMEM supplemented with 10% FBS and 1% Penicillin/Streptomycin. AML-12 cells were cultured in DMEM:F12 supplemented with 10% FBS, 1x ITS (insulin, transferrin, selenium), and 40 ng/ml dexamethasone. Freestyle 293-F cells (Gibco) were cultured in Freestyle 293 Expression Medium (Gibco) supplemented with 0.5% Penicillin/Streptomycin. C57BL/6 Mouse Primary Brain Microvascular Endothelial Cells were purchased from Cell Biologics and cultured in Complete Endothelial Cell Medium with the growth factor supplement (M1168, Cell Biologics). Primary T cells were cultured in Advanced RPMI 1640 supplemented with 10% FBS, 1% penicillin/streptomycin, 30U/ml of IL-2 (PeproTech), and 2.5 µl of Dynabeads Mouse T-Activator CD3/CD28 per 1×10^5^ cells.

### LNP Transfection and Fluorescence/Luminescence Readouts

Target cells were seeded as 20k/well on 96-well plates, the day before the transfection. For transfection, LNPs were incubated with the conditioned media or purified proteins for 15 minutes at 37°C. Post-incubation, they were dropped on cells and incubated for 24 hours. The uptake efficiency of GFP mRNA-encapsulated LNPs were assessed using fluorescence microscopy and flow cytometry. For fLuc mRNA-encapsulated LNPs, cells were treated with luciferin substrate (Bright-Glo Luciferase Assay System, Promega) according to manufacturer’s instructions, and luminescence were measured using a plate reader (BioTek Synergy).

### Constructs

ApoE WT and mutants, ApoE-ligand fusions, N-terminus secretion tag, haPCSK9 (hyperactive)…

### Secreted Protein Collection and Protein Purification

For secreted protein collection, HEK293FT cells were seeded on 24-well plates with 1.2×10^5^ cells/well density and transfected with the cloned constructs the next day. After overnight incubation, the media were changed to serum-free DMEM. After 72 hours of transfection, media containing the expressed and secreted proteins were collected. Expression and secretion of each construct were monitored by running samples on stain-free protein gels (Bio-Rad).

For large scale production and purification of proteins, Freestyle 293-F suspension cells were used. 200 million 293-F cells were seeded in 200 ml of FreeStyle 293 Expression Medium in a 1L flask on an orbital shaker in an incubator. The next day, cells were transfected with 200 µg of expression plasmids using 600 µl of PEI per flask. Cells were centrifuged after 3 days and media were filtered through 0.22 µm vacuum filters. Media were equilibrated with PBS containing 10 mM of imidazole (Teknova), in a 1:1 volume ratio and incubated with 3-4 ml of HisPur Ni-NTA Resin (Thermo Scientific) at 4°C for 1h. They were centrifuged for 2 min at 800g, and supernatants were discarded. Resins were resuspended with the equilibration buffer and loaded on Pierce Centrifuge Columns (Thermo Scientific). After centrifuging for 1 min at 500g, they were washed with PBS with 25 mM of imidazole for 5 times and eluted with PBS containing 250 mM of imidazole as 5 fractions. Each fraction was run on stain-free gels and the ones having purified protein samples were collected. Buffer exchange and concentration was performed with Amicon Ultra Centrifugal Filters (Millipore Sigma). Proteins were stored in PBS containing 1x Protein Stabilizing Cocktail (Thermo Scientific) at −20°C.

### Mass Spectrometry and Competition Assay

Protein A/G magnetic beads (APExBIO) were washed 3 times with PBS and incubated with 5 µg of anti-PEG antibody (ACROBiosystems) per 10 µL of beads for each sample, for 30 min at room temperature with gentle mixing. After washing 3 times with PBS to remove unbound antibody, LNPs (4 µg of RNA) were incubated with the antibody-coated beads in PBS with a final volume of 200 µL, for 15-20 min at room temperature with gentle agitation. Beads were washed 3 times with PBS to remove unbound LNPs. The bead-LNP complexes were incubated at 37 °C for 30 min with either 10% mouse serum, recombinant mutant ApoE (100 µg), or both depending on the experimental group in 200 µL final volume. Following incubation, beads were washed 5 times with 50 mM ammonium bicarbonate (ABC) buffer to remove unbound proteins. Protein coronas were eluted by incubating the beads with 100 µL of 0.1 M glycine-HCl buffer (pH 2.8) for 5 min at room temperature with intermittent vortexing. Supernatants were collected and immediately neutralized with 10-15 µL of 1 M Tris-HCl (pH 8.5). Samples were stored at 4°C overnight prior to mass spectrometry analysis. Eluted and neutralized proteins were submitted to the Taplin Mass Spectrometry Facility (Harvard Medical School) in low-binding tubes in 100 µL final volume. Downstream tryptic digestion and peptide cleanup were performed by the facility.

### Animal Experiments

All animal procedures were conducted in accordance with protocols approved by the Institutional Animal Care and Use Committees (IACUC) at the Massachusetts Institute of Technology (protocol #220800415) and Brigham and Women’s Hospital (protocol #2023N000166).

Control or experimental LNPs were administered to C57BL/6J mice by retro-orbital injection at doses ranging from 0.5 to 2 mg/kg mRNA. Six hours after injection, bioluminescence imaging was performed using an IVIS Spectrum system (Xenogen IVIS-200, MIT) or a Bruker MI Xtreme (BWH). D-luciferin potassium salt (Revvity) was prepared in PBS at 15 mg/mL and administered intraperitoneally at 150 mg/kg. After 5 minutes, mice were euthanized, and organs were harvested, rinsed in PBS, and immediately imaged using identical exposure settings. Luminescence intensity was quantified for each organ using the corresponding imaging software. Organs designated for downstream cell isolation were transferred to centrifuge tubes containing PBS supplemented with 2% FBS and kept on ice until processing.

### Isolation of Primary Cells

Primary T cells and Hematopoietic Progenitor Cells (HPC) were isolated using EasySep Mouse T Cell Isolation Kit (Stemcell Technologies) and EasySep Mouse Hematopoietic Progenitor Cell Isolation Kit, respectively (Stemcell Technologies) according to the manufacturer’s instructions. Briefly, spleens or bones were harvested and placed into a 1.5 ml centrifuge tube in the primary cell buffer (PBS containing 2% FBS and 1 mM EDTA). Tissues were chopped/crushed into small pieces using scissors to release cells into the solution. Debris was removed by passing cell suspension through a 70 µm mesh strainer (Greiner Bio-One). Cells were centrifuged at 300g for 10 min and resuspended at 1×10^8^ cells/ml in the primary cell buffer. 20 µl of FcR blocker were added and mixed. Samples were placed into a 14-ml round bottom tube, and 50 µl of T cell or HPC isolation cocktails were added. After 10 min of incubation at RT for T cells, or 15 min at 4°C for HPCs, 75 µl of RapidSpheres were added to samples. T cells were incubated for 2.5 min at RT; HPCs were incubated for 10 min at 4°C. Samples were topped up to 2.5 ml with a primary cell buffer, and placed into a magnet. After 5 min of incubation at RT, cell suspension cleared from spheres were pipetted into a 15-ml centrifuge tube.

Primary hepatocytes were isolated from liver tissue using a collagenase-based enzymatic digestion method. Briefly, excised liver tissue was rinsed in cold HBSS to remove residual blood. The tissue was finely minced using scissors. The minced tissue was transferred into a digestion solution containing 0.25 mg/ml Collagenase D and DNase I (50 μg/ml) in HBSS. The tissue suspension was incubated at 37°C for 60 minutes with gentle agitation in a rotating incubator. Enzymatic digestion was terminated by adding bovine serum albumin (BSA) to a final concentration of 1%, and the suspension was immediately placed on ice. The digested mixture was filtered through a 70 μm cell strainer to remove undigested debris. The filtrate was transferred into conical tubes and placed on ice for 5 minutes to allow hepatocytes to sediment by gravity. The supernatant, containing mostly non-parenchymal cells, was carefully discarded. The cell pellet was resuspended in cold PBS containing calcium and magnesium, and the sedimentation step was repeated twice to enrich hepatocytes.

### Flow Cytometric Detection of CAR Expression

For flow cytometric analysis of CAR expression, cells were stained using a PE-labeled monoclonal anti-FMC63 scFv antibody (Mouse IgG1, clone Y45; Acrobiosystems, Cat. No. FM3-HPY53), according to the manufacturer’s instructions. Primary T cells were isolated from LNP-treated mice, washed once with FACS buffer (PBS containing 2% FBS and 1 mM EDTA), and counted to determine viable cell numbers. An aliquot of 5 × 10^5^ viable cells was transferred into each tube.

The antibody stock solution was diluted 1:50 in the FACS buffer immediately before use (2 μl antibody per 5 × 10^5^ cells). A total of 100 μl of the diluted antibody solution was added to each cell pellet, mixed gently, and incubated for 60 min at 4°C. Following incubation, cells were washed three times with FACS buffer and resuspended in 200 μl PBS. Samples were analyzed for PE positivity using a flow cytometer (CytoFlex LX, Beckman Coulter).

### *In vitro* Cytotoxicity Assay

HEK293FT cells were seeded in 96-well plates (2×10^4^ cells/well). The next day, they were transiently transfected with human CD19 expression plasmid (Addgene #196634) using Lipofectamine 3000 (Thermo Fisher Scientific) according to the manufacturer’s protocol, 24 hours prior to co-culture. Additionally, they were treated with LNPs formulated with firefly luciferase mRNA, 8 hours prior to co-culture. T cells were isolated from mice treated with LNPs formulated with anti-CD19 CAR mRNA at 6-8 hours post-injection. A portion of the isolated cells was stained with an anti-FMC63 antibody (Acro Biosystems) to assess CAR expression by flow cytometry. The remaining T cells were activated for 2h by culturing in T cell activation media; Advanced RPMI 1640 supplemented with 10% FBS, 1% penicillin/streptomycin, 30U/ml of IL-2, and 2.5 µl of Dynabeads Mouse T-Activator CD3/CD28 per 1×10^5^ cells.

Activated CAR T cells were added on target HEK-CD19-Luc cells at 5:1 effector-to-target (E:T) ratios. Co-cultures were incubated in a 1-to-1 mix of regular HEK293FT media and T cell activation media, for 24 hours. Luciferase activity was measured using the Bright-Glo Luciferase Assay System (Promega) according to the manufacturer’s instructions. Luminescence was recorded using a plate reader (BioTek Synergy), and specific cytotoxicity was calculated by the percent decrease in the luminescence signal compared to control wells.

### *In vivo* partial rejuvenation of aged T cells

14-month-old mice received a single intravenous dose (1 mg/kg RNA) of LNPs encapsulating a pooled set of five mature microRNA mimics corresponding to the miR-302a/b/c/d and miR-367 cluster. LNPs were conjugated with a CD5 antibody to enable T cell targeting, and mice were pretreated with haPCSK9 15 minutes prior to LNP administration to reduce hepatic uptake. At day 1 post-injection, splenic T cells were isolated and a subset was immediately analyzed for DNA damage by γH2AX staining. The remaining cells were cultured ex vivo and analyzed again at day 5 and day 12 to assess longitudinal changes in DNA damage markers following a single in vivo treatment.

### Determining γ-H2AX Levels via Flow Cytometry

H2AX phosphorylation was examined using flow cytometry analysis following prior descriptions, with minor changes implemented. After in vivo treatment, harvesting, and culturing, the cells were counted and subsequently pelleted (4 min at 400g), then fixed in BD Phosflow™ Fix Buffer (4.2% formaldehyde) for a duration of 10 min at room temperature. Next, cells were permeabilized using BD Phosflow™ Perm/Wash Buffer I for 15 minutes. A final staining mixture was prepared at a 1:100 ratio: 1ul of Phospho-Histone H2A.X (Ser139, Monoclonal Antibody, CR55T33, PerCP-eFluor™ 710, eBioscience™) per 100ul of BD perm/wash buffer was mixed with cells and incubated in darkness for 45min at ambient temperature, with occasional agitation. The samples were subsequently washed with BD perm/wash buffer before being resuspended in BD Stain Buffer (FBS). These samples were analyzed on a Beckman Coulter Cytoflex LX utilizing the subsequent settings: PerCP-eFluor™ 710 was stimulated by the 561 nm Yellow-Green laser, and its fluorescence was acquired in the Y710 channel via a 710/50 nm bandpass filter. No compensation was necessary. Gating Strategy: The total T-cell population was identified on the SSC-A, FSC-A plots. Single cells were chosen, and then the mean γ-H2AX fluorescence signal was acquired as the ultimate measurement. This mean γ-H2AX fluorescence signal was subsequently transformed into a fold change metric normalized against control values.

### Statistics and reproducibility

All experiments were performed with at least three independent biological replicates unless otherwise stated. Statistical analyses and data visualization were conducted using GraphPad Prism v.10 (GraphPad Software LLC), and figures were assembled using Adobe Illustrator (Adobe Inc.). For comparisons between two groups, two-tailed unpaired Student’s *t*-tests were used. For experiments involving multiple conditions, statistical analyses were performed using appropriate pairwise comparisons relative to defined control groups, as indicated in the corresponding figure legends. Data are presented as mean ± SEM, and *P* < 0.05 was considered statistically significant. Experimenters were not blinded to group allocation, and sample sizes were not predetermined. Some Extended Data experiments were exploratory pilot studies and were performed once, as noted in the respective figure legends.

## Extended Data Figures

**Extended Data Fig. 1:**
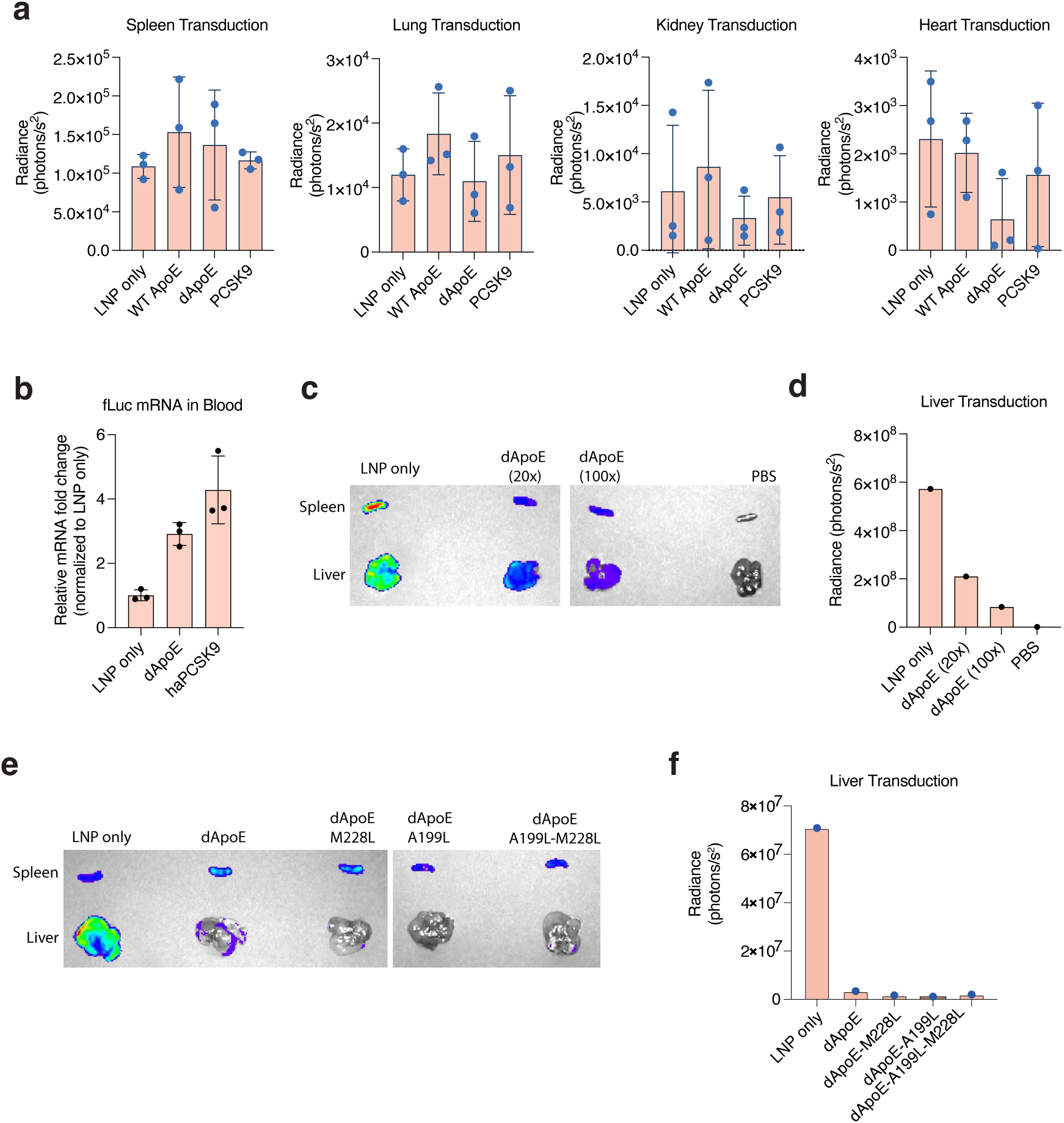
Dose dependence and lipid-binding domain variants of dApoE enable liver detargeting without altering extrahepatic biodistribution. (a) Ex vivo bioluminescence analysis showed that dApoE precoating or haPCSK9 pretreatment selectively reduced hepatic transduction while leaving signals in spleen, lung, kidney, and heart largely unchanged, indicating that liver detargeting does not cause nonspecific redistribution to other organs. (b) Consistent with reduced hepatic sequestration, circulating fLuc mRNA levels measured by qPCR were increased in blood following dApoE or haPCSK9 treatment, reflecting prolonged systemic persistence of LNPs due to decreased liver uptake. (c-d) Liver detargeting by dApoE was dose-dependent, as increasing dApoE:LNP ratios progressively suppressed hepatic bioluminescence. (e-f) Incorporation of selected lipid-binding domain mutations into dApoE further reduced liver transduction in vivo, suggesting that modulation of LNP-ApoE interactions can enhance detargeting efficiency. dApoE:LNP mRNA ratio is 100× for each variant.

**Extended Data Fig. 2:**
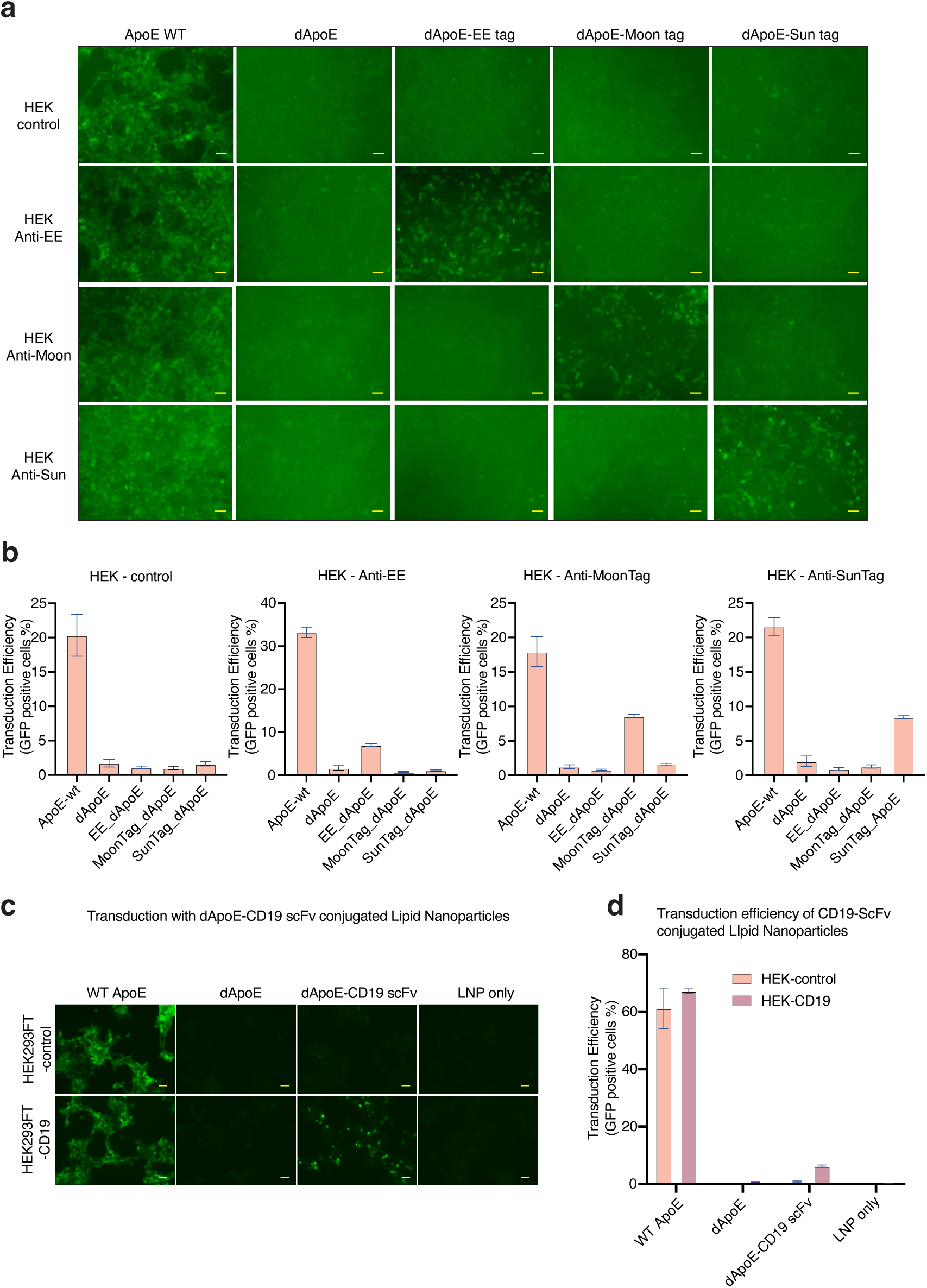
Retargeting via fusion constructs and synthetic binders. (a) Fluorescence imaging and (b) flow cytometry analysis of LNP uptake in HEK293FT cells expressing cognate surface binders demonstrate that synthetic binder-dApoE fusion constructs enable receptor-specific transduction only in engineered cell lines, with minimal background in control cells. Scale bar, 100 µm. (c-d) scFv-based retargeting was evaluated using a CD19-specific scFv genetically fused to dApoE. While dApoE-CD19 scFv fusion LNPs mediated selective uptake in CD19-overexpressing HEK293FT cells relative to control cells, the magnitude of enhancement was modest and substantially lower than that achieved with wild-type ApoE.

**Extended Data Fig. 3.**
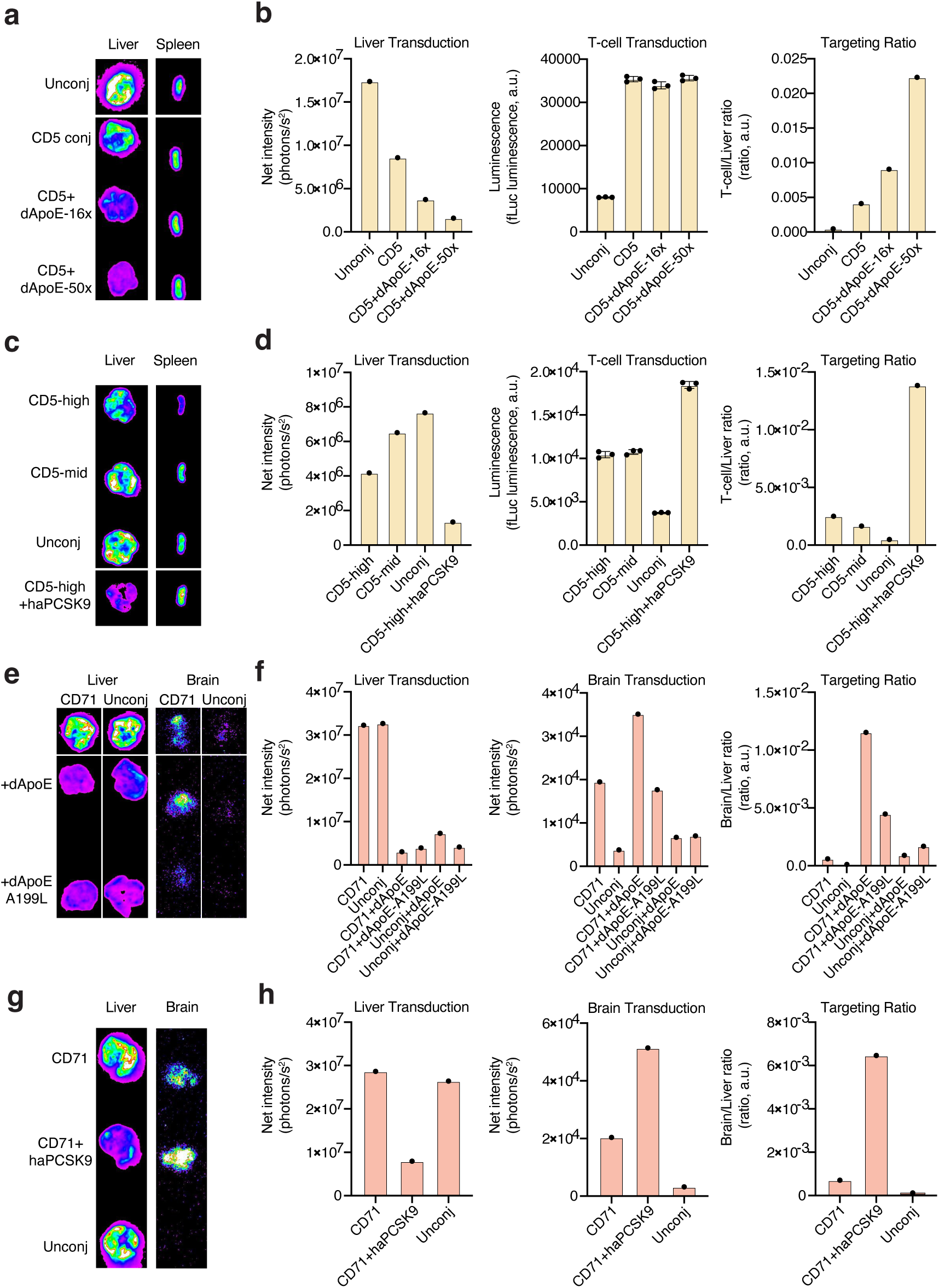
Pilot optimization of antibody conjugation and detargeting parameters for T cell and brain targeting in vivo. (a-b) CD5 antibody–conjugated LNPs were evaluated with increasing dApoE:LNP mRNA ratios (16× and 50×), revealing progressive suppression of hepatic transduction while preserving T cell delivery and substantially improving T cell-to-liver targeting ratios. (c-d) Antibody density was further optimized by comparing CD5-LNP conjugation ratios, with “CD5-high” and “CD5-mid” corresponding to antibody:mRNA ratios of 2.2 and 1.1, respectively; increasing antibody density enhanced T cell targeting ratio which was further enhanced by haPCSK9 pretreatment. (e-f) For brain targeting, CD71-conjugated LNPs were combined with either dApoE (30×) or the lipid-binding–domain variant dApoE-A199L (15×), resulting in reduced hepatic signal and improved brain-to-liver ratios compared to unconjugated or non-detargeted controls. dApoE provided a better brain targeting ratio. (g-h) At an increased LNP dose (0.75 mg/kg) and higher haPCSK9 dosing (60 µg per mouse), haPCSK9 pretreatment further enhanced brain targeting by CD71-conjugated LNPs, increasing brain transduction while maintaining strong suppression of liver uptake.

**Extended Data Fig. 4.**
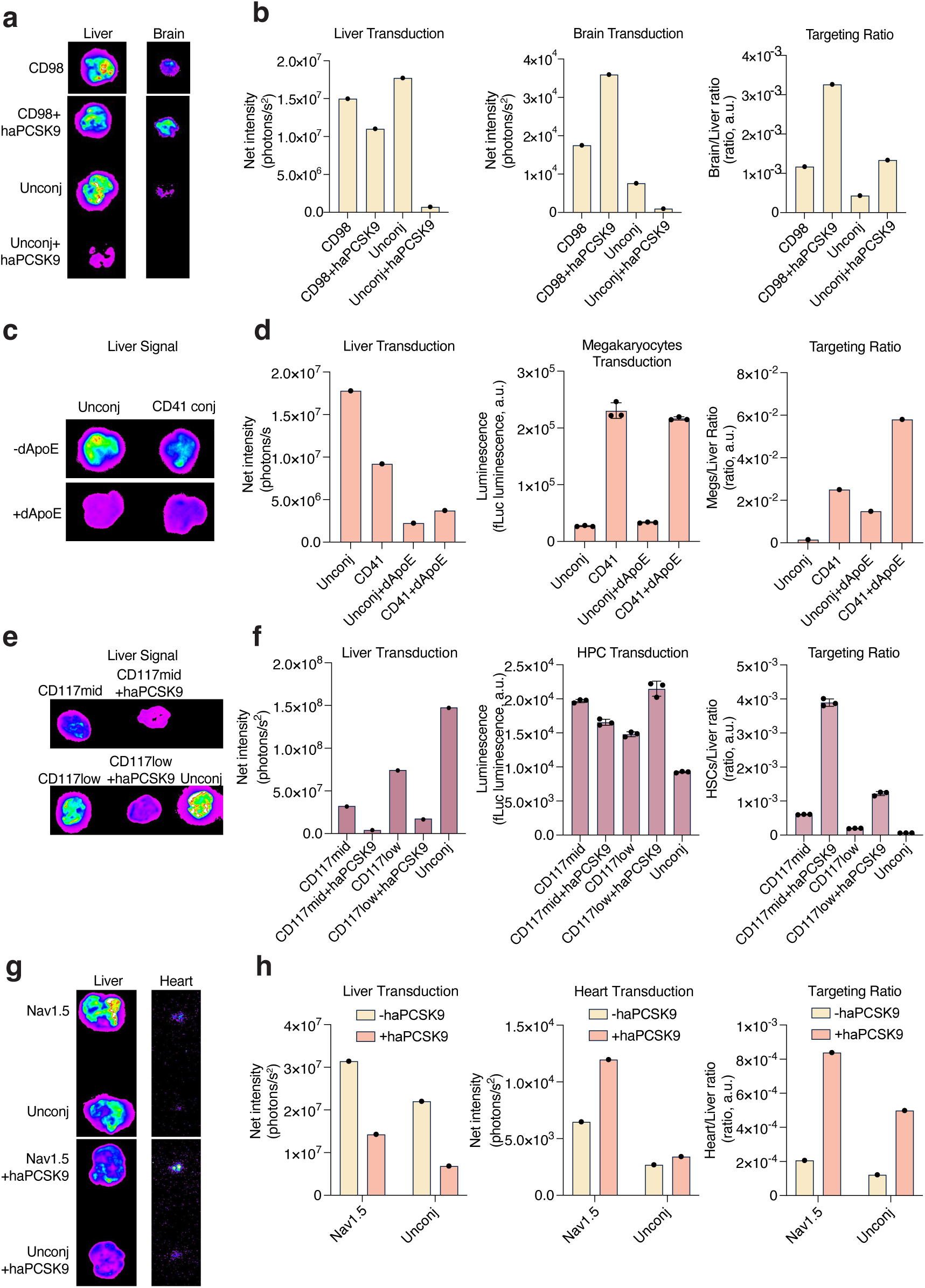
Pilot targeting of additional cell types using antibody-conjugated LNPs combined with detargeting strategies. (a-b) CD98 antibody-conjugated LNPs were evaluated for brain-associated delivery at an LNP dose of 0.5 mg/kg with an antibody:mRNA ratio of 0.5; haPCSK9 pretreatment (40 µg per mouse) reduced hepatic transduction while increasing brain-to-liver targeting ratios relative to unconjugated controls. (c-d) Megakaryocyte targeting was assessed using CD41 antibody-conjugated LNPs (0.5 mg/kg, antibody:mRNA ratio 0.5) with or without dApoE precoating (30× relative to LNP mRNA), showing strong megakaryocyte transduction and marked suppression of liver signal in the presence of dApoE. (e-f) Hematopoietic progenitor cell (HPC) targeting was examined using CD117 antibody conjugation at two antibody densities (mid ratio 1.1 and low ratio 0.5) combined with haPCSK9 pretreatment (40 µg per mouse, LNP dose 0.5 mg/kg), resulting in reduced liver uptake and preserved or enhanced HPC transduction depending on antibody density. (g-h) Cardiac targeting was explored using Nav1.5 (SCN5A) antibody-conjugated LNPs (0.5 mg/kg, antibody:mRNA ratio 0.5), where haPCSK9 pretreatment further suppressed hepatic background and increased heart-to-liver targeting ratios. Together, these pilot experiments demonstrate that antibody conjugation enables delivery to diverse target cell types, while detargeting strategies consistently reduce hepatic uptake and can improve apparent targeting specificity across multiple tissues.

**Extended Data Fig. 5.**
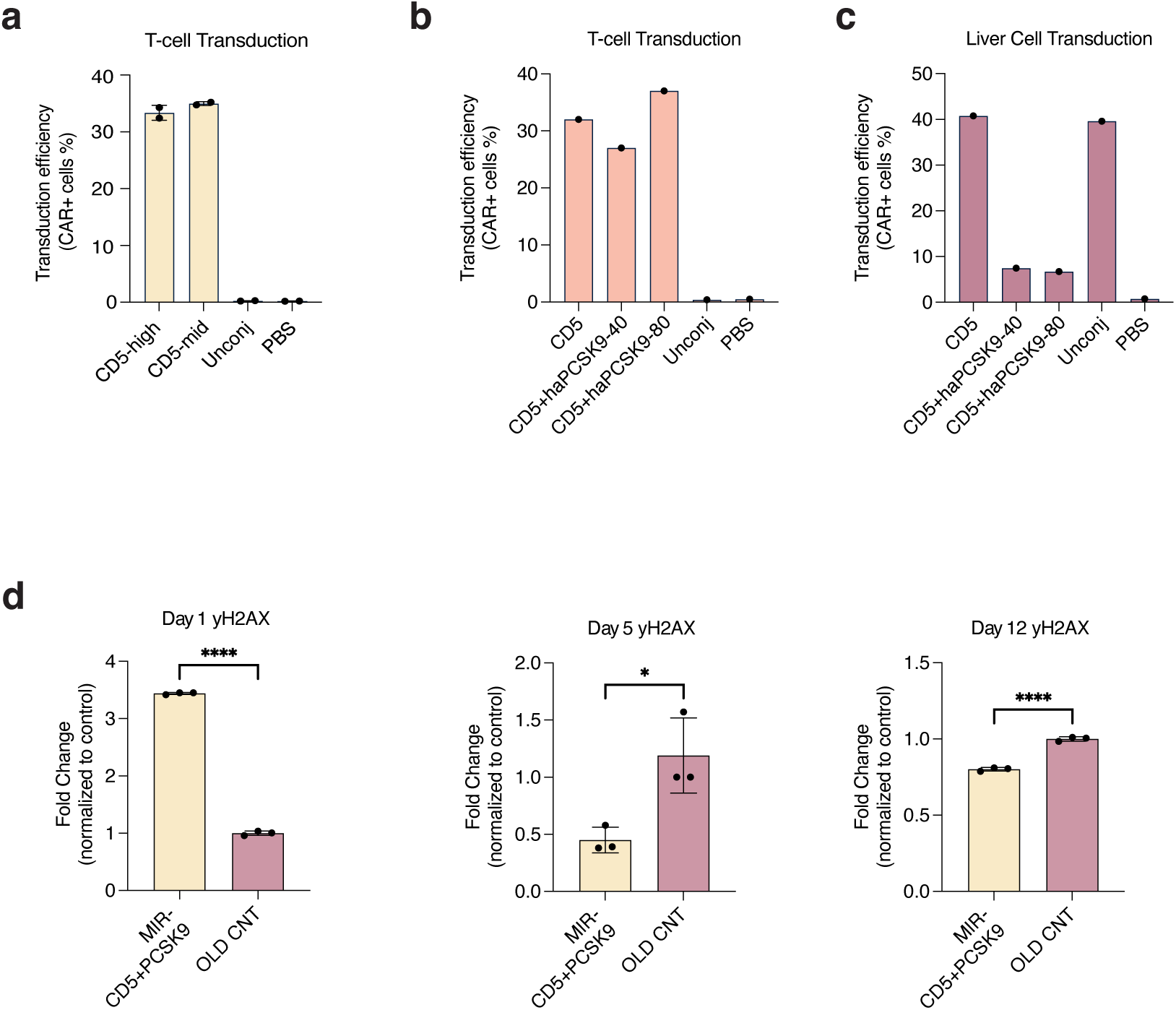
Pilot optimization of in vivo CAR T cell generation and targeted delivery of partial reprogramming signals to aged T cells. (a) Flow cytometry-based quantification of CAR+ T cell frequencies following systemic delivery of CD5 antibody-conjugated LNPs demonstrates the effect of antibody density at an increased LNP dose (1 mg/kg RNA). “CD5-high” and “CD5-mid” correspond to antibody:mRNA ratios of 2.2 and 1.1, respectively, and yielded comparable CAR+ T cell frequencies (∼30-35%), indicating that antibody density was not a major determinant of efficiency under these conditions. (b) Using the CD5-mid conjugation ratio, CAR+ T cell frequencies were measured after haPCSK9 pretreatment at 40 µg or 80 µg per mouse, revealing a modest improvement in apparent T cell targeting with the higher haPCSK9 dose. (c) In parallel, flow cytometric analysis of liver cells showed that both haPCSK9 doses comparably suppressed off-target hepatic transduction. (d) CD5 antibody-conjugated LNPs delivering a pool of miR-302a/b/c/d and miR-367 mimics (1 mg/kg RNA) were administered following haPCSK9 pretreatment. γH2AX levels in isolated T cells were quantified at day 1, day 5, and day 12 post-injection and normalized to age-matched untreated controls. An early increase in γH2AX was observed at day 1, followed by a significant reduction at days 5 and12. Data are shown as mean ± SEM.

## Acknowledgements

We thank all members of Abudayyeh-Gootenberg Lab for support and advice. J.S.G. and O.O.A. are supported by NIH grants 1R21-AI149694, R01-EB031957, R01-AG074932 and R56-HG011857; Rett Syndrome Research Trust; Gordon and Betty Moore Foundation; Evolution and AFAR; Google Deepmind; Human Immunome Project and Michelson Medical Research Foundation; the G. Harold & Leila Y. Mathers Charitable Foundation; the NHGRI Technology Development Coordinating Center Opportunity Fund; the MIT John W. Jarve (1978) Seed Fund for Science Innovation; Impetus Grants; a Cystic Fibrosis Foundation pioneer grant; Google Ventures; FastGrants; the Harvey Family Foundation; Winston Fu; and the McGovern Institute.

## Contributions

A.K. and C.S. conceived the study and participated in the design, execution and analysis of experiments. A.K., C.S., A.E., F.F., K.D., and S.S. participated in the design and execution of in vivo trials. A.E., F.F., K.D., A.X.N, I.H, A.A., and S.P.N. assisted with readouts, generated lipid nanoparticles, performed tissue culture, purified protein, and generated constructs for evaluation. A.A., K.D., A.X.N, I.H., A.E., and F.F. purified mammalian protein. P.T.P. and S.L. participated in design and execution of in vivo reprogramming experiments. C.F. assisted with analysis and readout of experiments. J.S.G, O.A., A.K., and C.S wrote the manuscript. A.K. and C.S. share co-first authorship.

## Ethics Declarations

Competing interests: Brigham and Women’s Hospital Inc, Beth Israel Deaconess Medical Center Inc, and Massachusetts Institute of Technology have filed for a patent application for this work (WO2025072956A2 and WO2025072956A3). J.S.G. and O.O.A. are co-founders of Terrain Biosciences, Monet Therapeutics, and Transit Therapeutics. All other authors declare no competing interests.

## Data availability

Proteomics data have been deposited at PRIDE. Expression plasmids are available from Addgene under the UBMTA; support information is available at https://www.abugootlab.org/. All other data are available from the corresponding authors upon reasonable request.

